# Covert detection of own-name and semantic violations in task-irrelevant speech, in a realistic Virtual Café

**DOI:** 10.1101/2022.07.06.498989

**Authors:** Adi Brown, Danna Pinto, Ksenia Burgart, Yair Zvilichovsky, Elana Zion-Golumbic

**Author notes:** Corresponding Author Prof. Elana Zion Golumbic, The Gonda Center for Multidisciplinary Brain Research Bar Ilan University, Ramat Gan, Israel, www.golumbiclab.org. joint first authors (Adi Brown, Danna Pinto).

## Abstract

Detecting that someone has said your name is one of the most famous examples for incidental processing of supposedly-unattended speech. However, empirical investigation of this so-called “cocktail party effect” has yielded conflicting results. We present a novel empirical approach for revisiting this effect under highly ecological conditions, by immersing participants in a multisensory virtual café environment and using realistic stimuli and tasks. Participants listened to conversational speech from a character sitting across from them, while a barista in the back of the café called out food orders. Unbeknownst to them, the barista sometimes called orders containing their own name or semantic violations. We used combined measurements of brain activity (EEG), eye-gaze and galvanic skin response to assess the response-profile to these two probes in the task-irrelevant barista-stream.

Both probes elicited unique neural and physiological responses relative to control stimuli, indicating that the system indeed processed these words and detected their unique status, despite being task-irrelevant. Interestingly, these responses were covert in nature and were not accompanied by gaze-shifts towards the barista character. This pattern demonstrates that under these highly ecological conditions, listeners incidentally pick up information from task-irrelevant speech, emphasizing the dynamic and non-binary nature of attention in real-life environments.

## Introduction

The cognitive and neural mechanisms that enable people to successfully focus on one speaker of interest in multi-talker environments have been a matter of ongoing interest for decades (Cherry, 1953; Bronkhorst, 2015; Qian et al., 2018). A major point of theoretical debate has been the extent to which speech that is irrelevant to the listener, and which they supposedly try to ignore, is nonetheless encoded and processed. Sparking this debate are conflicting behavioral and neural results. Many studies report evidence for encoding of task-irrelevant speech only at a sensory level (Cherry, 1953; Näätänen et al., 1997; Escera et al., 2003; Paavilainen, 2013; Ding et al., 2018; Parmentier et al., 2019), in line with early selection models of attention (Broadbent, 1965). However, others show evidence that listeners can extract some linguistic and/or semantic attributes from task-irrelevant speech (Moray, 1959; Lachter et al., 2004; Bronkhorst, 2015; Röer et al., 2017; Brodbeck et al., 2020; Agmon et al., 2021), which is more in line with late-selection and attenuation models of attention (Deutsch and Deutsch, 1963; Treisman, 1964). These include priming and intrusion effects of irrelevant speech (Dupoux et al., 2003; Rivenez et al., 2008), neural response to some linguistic and semantic features (Brodbeck et al., 2020; Agmon et al., 2021; Holtze et al., 2021), as well as explicit detection of salient words such as one’s own name (which is reported in 30%-96% of participants; Moray, 1959; Wood and Cowan, 1995; Conway, Cowan and Bunting, 2001; Tamura *et al*., 2012; Tateuchi, Itoh and Nakada, 2012; Naveh-Benjamin *et al*., 2014; Röer and Cowan, 2021) or semantically salient words (Van Petten and Luka, 2012; Röer et al., 2017).

Rather than expecting the answer to whether task-irrelevant speech is processed linguistically to be a dichotomous ‘yes’ or ‘no’, it is likely that the level of processing applied to task-irrelevant speech depends on the specific context in which this is tested (Lavie, 2005; Murphy et al., 2017). Unfortunately, the majority of research into this question has used extremely artificial paradigms and stimuli, that are far removed from the listening challenges of daily life, which probably contributes to the inconsistency of results across studies. For example, many dichotic listening experiments investigating the “irrelevant sound effect” use arbitrary lists of words as stimuli that lack contextual continuity (Lewis, 1970; Bentin et al., 1995; Wood and Cowan, 1995; Conway et al., 2001; Tun et al., 2002; Naveh-Benjamin et al., 2014; Vanthornhout et al., 2019). Moreover, in most of those studies, the tasks that participants were asked to perform are also extremely artificial and vary in their difficulty, such as committing word-lists to short-term memory or detecting a prescribed target word. Hence, even though these studies offer important insights into some aspects of the competition for linguistic processing between concurrent speech, they leave much to be desired if we strive to understand how irrelevant speech is processed under naturalistic conditions. In other words, the well-known ‘cocktail party effect’ has actually not been tested in a true to life cocktail party setting.

The goal of the current study was to investigate this long-standing question under ecological conditions, that simulate the type of speech-stimuli, tasks and environment that a listener may encounter in real-life. To this end, we created a Virtual Café where participants experience conversing with a partner in a realistic café environment and hear speech from additional characters in the café (Shavit-Cohen and Zion Golumbic, 2019). Participants sat across from an animated avatar character (‘target speaker’) and were told to listen to their narrative of conversational speech. A barista-character was presented in the back of the café, calling out orders (e.g., “Salad for Sarah”), however this stream was task-irrelevant. Critically, we manipulated the content of this task-irrelevant stream in two ways: (1) on some occasions, the orders called contained the participants’ own name; or (2) the orders contained a semantic violation (e.g., “Coffee for salad”). This unique setup allowed us to revisit two central hypotheses regarding the special affinity for detecting one’s own name (Moray, 1959; Wood and Cowan, 1995; Holtze et al., 2021) and detecting semantic properties of task-irrelevant speech (Van Petten and Luka, 2012), and to do so under ecologically relevant conditions, where these stimuli might actually occur in real-life. We measured participants’ neural activity, gaze-dynamics and galvanic-skin-response (GSR) as they engaged in listening to their conversation partner in the virtual café. This multi-dimensional data set allowed us to assess the variety of ways in which listeners may (or may not) respond to semantically or personally salient events in task-irrelevant speech, under these ecologically relevant conditions.

## Materials and Methods

### Participants

Fifty adults participated in this study (ages 19-34, median 23; 32 female, 18 male, ten left-handed), all fluent in Hebrew, with self-reported good eyesight or corrected by contact lenses, normal hearing, and no history of neurological disorders (9 reported being diagnosed with ADD/ADHD). Due to technical issues and excessive artifacts, some participants were excluded from the analyses of different metrics (eye-gaze, neural, GSR), as reported below for each measure. Signed informed consent was obtained from each participant prior to the experiment, in accordance with the guidelines of the Institutional Ethics Committee at Bar-Ilan University. Participants were paid for participation or received class credit.

### Apparatus

Participants were seated in an acoustic-shielded room and viewed a 3D virtual-reality (VR) scene of a café, through a head-mounted device (Oculus Rift Development Kit 2). The virtual environment, which consisted of several avatars sitting at various locations within the café, was programmed and presented using the software Unity (Figure 1A). Two of the avatars (the target-speaker and the barista, see below) were speaking, with their speech-audio manipulated so it was perceived as originating from their location in virtual space (3D sound algorithm, Unity). The audio was presented through headphone (Etymotic ER-1), and both the graphic display and 3D sound were adapted on-line in accordance with head-position, to maintain a spatially coherent experience of the virtual space. The articulation movements of the speaking avatars were synchronized to the envelope of the speech, to create a realistic audio-visual experience. Avatar eye-movements were generated to appear natural. Additional animations of avatar body movements were selected from the Mixamo platform (Adobe Systems, San Jose, CA, USA), adapted to the scene and randomly looped to avoid noticeable exact repetitions.

**Figure 1.**
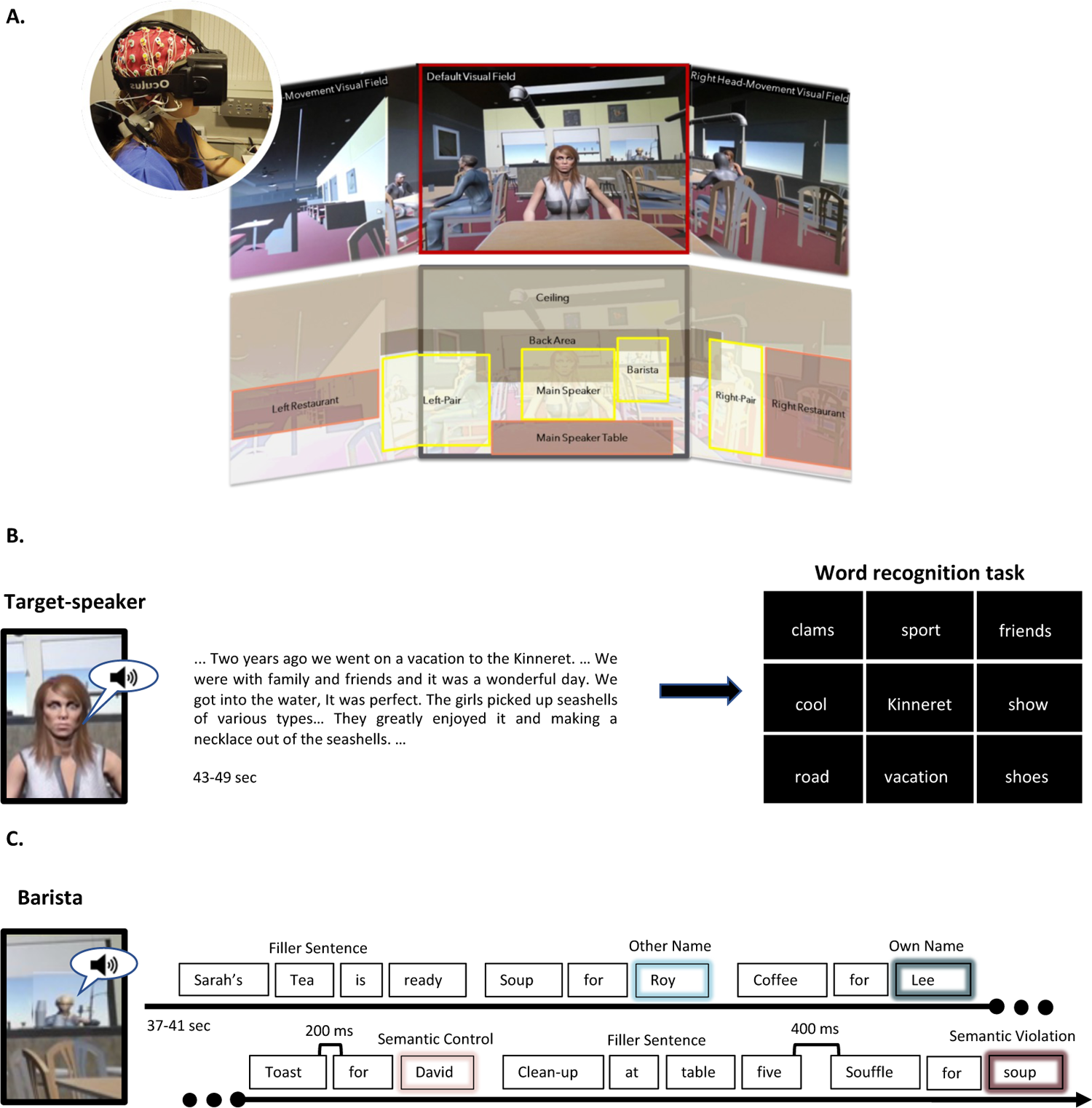
**A.** Illustration of the virtual café scenario. Top: A wide-view perspective of the café, as seen by the participants. The central area mark in red is the default visual field, whereas seeing the areas to the left and right require head-movements. Bottom: Demarcation of the regions-of-interest (ROIs) used for analyzing eye-gaze patterns. ROIs that contain moving avatars are marked in yellow. Circular inset: Participant wearing the Virtual-Reality headset over the EEG cap. **B.** Example of the narrative speech spoken by the target-speaker. Each trial was followed by a word recognition test, in which participants were asked to identify words that had been present in the target-speech, out of a 9 word-matrix. **C.** Illustration of the structure of the barista speech. This stimulus consisted of order-sentences, which contained probe-words (own-name and semantic violation) and their respective controls.

The VR device was custom-fitted with an embedded eye-tracker (SMI, Teltow, Germany; 60 Hz monocular sampling rate) for continuous monitoring of participants’ eye-gaze position. Gaze location was projected onto the virtual environment and was recorded as coordinates in three-dimensional space, as well as in terms of which object in the virtual environment participants looked at (e.g., target-avatar, different avatars, ceiling etc.).

Neural activity was recorded using a 64 electrodes EEG system (BioSemi; sampling rate: 1024 Hz) and a standard EEG cap. Two external electrodes were placed on the mastoids and served as reference channels. Electrooculographic (EOG) signals were simultaneously measured by 3 additional electrodes, located above the right eye and on the external side of both eyes. Figure 1A (circular inset) shows the placement of the VR head device over the EEG cap. Galvanic Skin Response (GSR) was also measured using two passive Nihon Kohden electrodes placed on the fingertips of the index and middle fingers of participants’ non-dominant hand. The signal was recorded through the BioSemi system amplifier and was synchronized to the sampling rate of the EEG.

### Stimuli

The virtual scene included the following avatar characters: a character sitting at a table facing the participant (target speaker), a barista standing at the bar in the back of the café, slightly to the right, and four additional characters sitting at tables to the left and right of the target speaker. The latter characters did not speak, however their body motions where slightly animated to preserve a realistic feel to the environment. The target-speaker and the barista were both presented as speaking.

The **target speaker’s speech** (Figure 1B) consisted of segments of natural Hebrew narratives pre-recorded by a female actor. To encourage spontaneous conversational-style speech, the actor was given a series of prompts and was asked to speak about them for approximately 40 seconds (timer shown on screen). The prompts referred primarily to daily experiences and could be of either a personal nature (e.g., “describe a happy childhood memory”) or more informational (e.g., “describe how to prepare your favorite meal”). The actor was given the list of prompts in advance in order to plan their answers, however the narratives themselves were delivered in a spontaneous, unscripted fashion. The amplitude of all narrative recordings was equated using the software PRAAT (Boersma and Weenink, 2010). In total, we used 32 narrative recordings, of which 2 were used for training, ranging between 43–49 seconds in length. Each narrative was presented only once during the experiment.

The **barista’s speech** (Figure 1C) consisted of lists of orders and filler sentences that might be heard in a café. The stimuli were constructed from single words, recorded individually by a female actor, in no systematic order. Recordings of individual words were cut and equated for amplitude (using PRAAT), and then concatenated (using Matlab) to create order-sentences, that consist of a person’s name and a food-item they had ordered. Order-sentences had two possible syntactic structures, which in English correspond to *“Salad for Sarah”*, or *“Sarah’s salad is ready”*. Importantly, in Hebrew, in both of these structures the name appears in the 3^rd^ word position of the sentence (“*Salat bishvil Sara*h” vs. “*hasalat shel Sarah mukhan*”), and hence its occurrence is similarly predictable. Besides order-sentences, additional filler sentences were constructed, that did not contain names and are contextually relevant for a café environment (e.g., *“Clean-up needed at table five”*, “*Lunch served until four”)*. In all sentences, words were presented with a constant between-word interval of 200 ms. Individual sentences were further concatenated into streams, with a 400 ms between-sentence interval to create streams of 37-41 seconds in length (each containing 13 sentences).

Some items in the barista-stream were manipulated, in accordance with our two research questions, serving as probe-words (Figure 1C): (1) **Own name** – some order-sentences contained the participant’s own name as the subject of the sentence. Own-names were presented in accordance with participants’ preferred pronunciation, based on verification during a prescreening phone call. Two potential control-names were chosen for each participant, which were the names of the two previous participants. These potential control-names occurred with equal probability as the own-name. In post-experiment debriefing, participants were asked about their familiarity with the two potential control-names, and the less familiar control-name was chosen for subsequent analyses. (2) **Semantic violation** – in some order-sentences a food-item was presented instead of the expected name (e.g., “*Coffee for salad*”), forming a semantically incongruent sentence. A total of 18 food item words were used as semantic-violation probe-words. These were matched with 18 control words – names presented in correctly structured sentences that occurred with similar probability and had similar syllable features (number; open or closed) and accentuation as the words used as semantic-violation probes.

For each participant, 30 personalized streams were constructed, each of which contained two probe-words of each type and their respective controls, as well as additional filler sentences. The sentences were concatenated in a pseudo-random order to ensure that sentences with own-name or semantic-violation probes were separated by at least two control sentences, and were not the first or last in the sequence.

### Procedure

Participants were instructed to listen to content of the target-speaker’s narrative and performed a word-recognition task following each trial. They were told that they might hear additional stimuli in the café, however that these stimuli were irrelevant. The word-recognition task following each trial consisted of presentation of a 9-word matrix, of which participants were asked to select all the words that they remember hearing in the target-speech. The number of correct words varied between 4-5, and participants received feedback on their responses after each trial (see calculation below). Proceeding to the next trial was controlled by the participants, and they were encouraged to take short breaks at regular intervals. Before the start of the experiment, participants were familiarized in two training trials with the virtual café scene, the task-relevant speaker and the task.

After completing the main task, participants were debriefed about their experience, and whether they noticed any stimuli that were out of the ordinary in the barista stream. Participants were also interviewed about their personal familiarity with people who share their own name or one of the two potential control-names (in order to select the control name that was least familiar).

After completing the main task, participants performed an additional task to assess their working memory capacity. We used a Hebrew adaptation (https://englelab.gatech.edu/translatedtasks.html#hebrew) of the Operation Span Working memory task (OSPAN), which is based on a Shortened Version of the working memory task by Foster *et al*. (2014). This task required the participants to solve simple arithmetic problems while also remembering a series of Hebrew letters. Each trial consisted of several (3-7) arithmetic problems, each followed by a single letter of the Hebrew alphabet. At the end of each trial the participants were asked to recall the series of letters in the order that they were presented. The test produces two main scores referring to the number of letters recalled and positioned correctly within the sequence (1) including only trials where all letters were correctly recalled (absolute OSPAN score) and (2) including all trials (partial OSPAN score). However, we used only the partial scores, as per recommendation of previous studies, since they make use of all available information (Friedman and Miyake, 2005; Redick et al., 2012; Đokić et al., 2018). 4 participants were removed from this analysis due to low performance (below 85% correct on the arithmetic problems; n=3) or due technical issues (n=1).

### Behavioral Data Analysis

The word recognition task consisted of detecting which of the 9 words were present in the target-speaker’s narrative. If participants detected all the words correctly and made no false-alarms, they received a score of 9/9 = 100%. This score was penalized by −1/9 (11%) if a correct word was missed *or* if an incorrect word was marked. Hence, this behavioral score reflects sensitivity to detecting the words from the narrative, combining both hits and correct-rejections.

### Eye-Gaze Analysis

The 3D café scene was parsed into several visual Regions of interest (ROI), as shown in Figure 1A. These included the areas around the characters in the café (*Main Speaker, Left-Pair, Right-Pair, Barista*) as well as regions of the café that did not contain characters (*Main Speaker Table, Ceiling, Floor, Bar-Left of Barista, Bar-Right of Barista, Back Wall, Left Restaurant, Right Restaurant*).

Information regarding participants’ momentary head-position and direction of gaze vector were combined to determine which ROI participants were looking at, on a moment to moment basis. Analysis of eye-gaze pattern focused on how long participants stayed fixated within a certain ROI as well as detection of saccades between ROIs, but saccades *within* ROIs were not analyzed, since this was outside the scope of our research interest.

First, we estimated global statistics of gaze-patterns, regardless of experimental manipulation. These included the percentage of each trial that a participant spent looking at each ROI and the rate of saccades between ROIs per trial. Then, we assessed whether probes elicited overt gaze-shifts by analyzing event-related gaze dynamics in 2-second-long time-windows following the probe words and their respective controls. In this analysis, we looked at the prevalence of saccades performed from the Main Speaker ROI to the Barista ROI, where the probe-words originated from. Due to technical issues with the eye-gaze data 10 participants were excluded from the analysis of global gaze-patterns, and 13 additional participants were excluded from the event-related analysis, yielding a total of n=40 and n=27 participants in each analysis, respectively.

### EEG Analysis

EEG preprocessing and analysis was performed using the matlab-based FieldTrip toolbox (Oostenveld et al., 2011) as well as custom written scripts. Raw data was first visually inspected and gross artifacts exceeding ±50 μV (that were not eye-movements) were removed. Independent Component Analysis (ICA) was performed to identify and remove components associated with horizonal or vertical eye movements as well as heartbeats. Any remaining noisy electrodes that exhibited either extreme high-frequency activity (>40Hz) or low-frequency activity/drifts (<1 Hz), were replaced with the weighted average of their neighbors using an interpolation procedure. Five participants were excluded from the EEG analysis due to extreme artifacts, and data is reported from the remaining n=45.

The main goal of this study was to investigate the neural response to words in the task-irrelevant barista-stream, and particularly to test whether the two type of probe words - hearing one’s own name or the presence of semantic violations - elicit unique neural responses, which would be indicative of their implicit detection. Therefore, our EEG analysis focused on the time period immediately after the presentation of each probe word and their respective controls. The clean EEG data was segmented into epochs between −200 to 1500 ms around each probe word and a 12Hz lowpass filter was applied.

Since the stimuli used as probe words were of varied lengths, ranging from 500-750 ms, and had different temporal profiles from each other, the classic method for deriving event-related potentials (ERPs) through simple averaging was not appropriate. Instead, we applied Residue Iteration Decomposition (RIDE) analysis (http://cns.hkbu.edu.hk/RIDE.htm) (Ouyang et al., 2011), which accounts for latency differences across trials when extracting the average neural response. Specifically, the RIDE analysis decomposes the data into stimulus-locked (S), and general (C) components, and analyzes the trial-to-trial variability of these components to reconstruct an event-related potential. The time-windows defined for the S and C components were 0-400 ms and 300-1200 ms respectively, values chosen based on visual inspection of the ERP grand average (prior to RIDE-estimation; Figure 7A inset, Figure 8A inset). In the supplementary material we report additional RIDE analyses conducted using slightly different time-window selections, however results are qualitatively similar. This analysis yielded a RIDE-ERP time course for each participant. These were then averaged across participants to derive a grand-average RIDE-ERP at each electrode. Statistical comparison of the responses to each probe-type (own-name and semantic-violations) vs. their controls was conducted by applying a t-test at each time-point between 200-1000ms and correcting for multiple comparisons (cluster permutation test, with threshold-free cluster enhancement correction (TFCE); implemented in the fieldtrip toolbox).

### GSR Analysis

Analysis of the GSR signal was performed using the matlab-based Ledalab toolbox (Benedek and Kaernbach, 2010), as well as custom written scripts. The raw data was manually inspected for distinguishable artifacts, which were fixed using spline interpolation. Then a continuous decomposition analysis (CDA) was performed on the full GSR signal and the time-locked response to individual probe words was extracted in windows between 0 and 5 seconds from the onset of each probe word. GSR responses to each probe word were averaged across trials, and a group-level grand-average was derived. Statistical analysis of the difference in responses to each probe-word vs. its control was conducted by applying a t-test to the mean amplitude within a 2-seconds time window surrounding the peak.

## Results

### Behavioral task

Word-recognition performance was relatively good, with an average accuracy of 85.27 ± 3.42% (std; Figure 2A), indicating that for the most part, participants paid attention to the target-speaker and correctly recognized words from the narrative. Note that this measure of accuracy is affected both by hits and correct rejections, reflecting sensitivity. Performance on the task was not correlated with working memory capacity (OSPAN partial; n=46, R^2^=0.11, p=0.49; Figure 2B).

**Figure 2:**
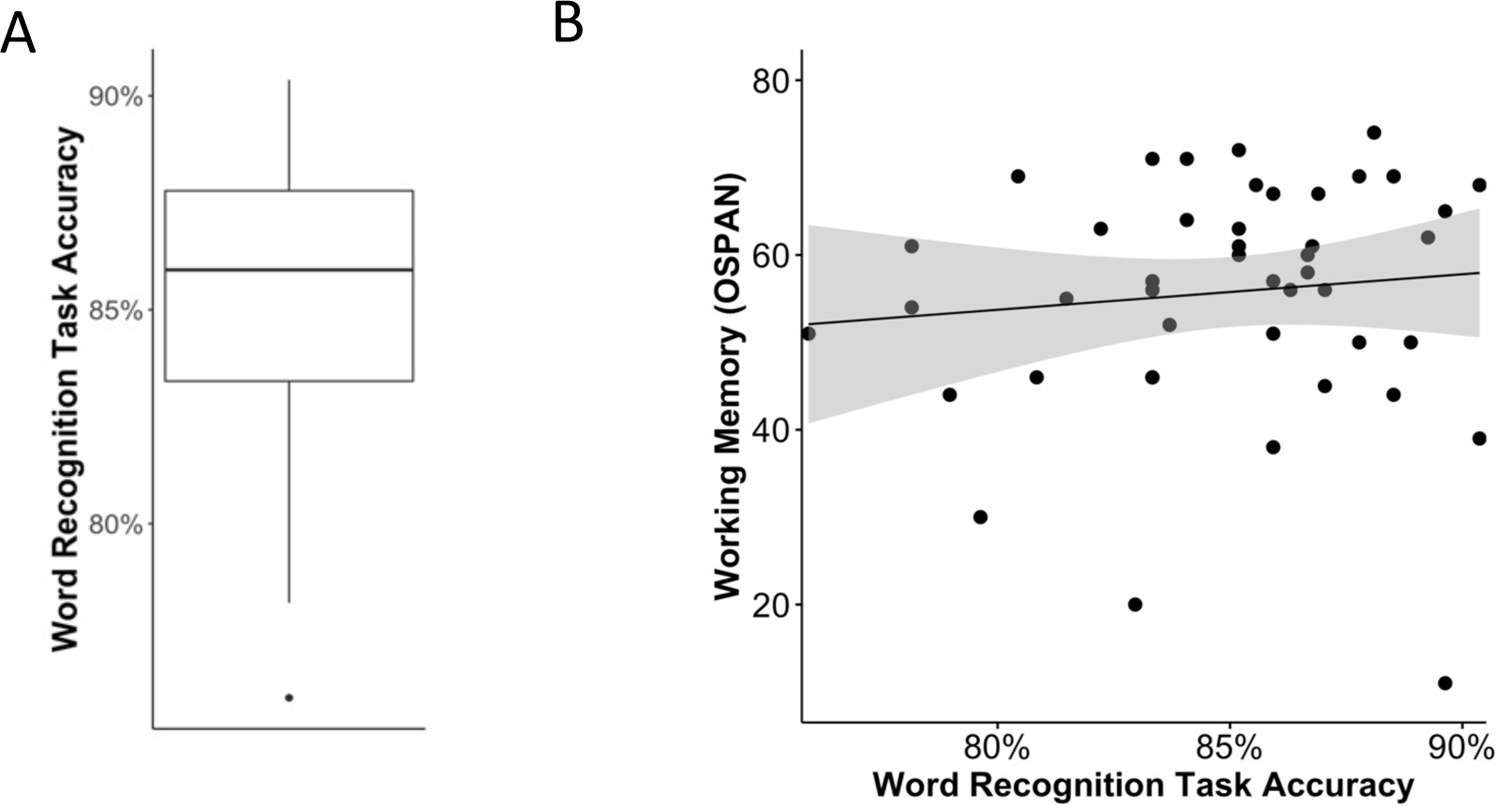
**A**. Box plot depicting the mean behavioral performance on the word recognition task (for all the participants before any exclusions, n=50). Error bar indicates SEM. **B**. Scatter plot of accuracy on the word recognition task vs. working memory capacity (OSPAN score), showing no significant correlation between them. The grey shaded area represents a 95% confidence interval. 4 participants were excluded from this analysis due to lack of a reliable OSPAN score (n=46).

### Post-experiment debriefing

After completing the experiment, participants were interviewed about their experience. In response to the question “how was the experiment?” 94% voluntarily mentioned that they had heard their name in the barista-stream, and 4% mentioned hearing sentences that did not make sense. Then, when asked directly “did you hear your name”, all but one participant recalled that they may have heard their own name or a similar name. When asked directly “did everything you heard make sense?” 4% responded that they noticed some portions of the speech that did not make sense, and additional 10% reported they felt like something wasn’t right, but could not put a finger on it, as they “didn’t really listen to the background”.

### Eye Tracking results

We first present a qualitative analysis of global gaze-patterns, regardless of the experimental manipulation (Figures 3-5). Although this analysis does not directly address to our research questions, it provides important insight into what participants actually did while experiencing the virtual café and how they naturally allocated their overt attention over time, in this rich multisensory scene. We then present the event-related analysis, aimed at determining whether the two probe-words placed in the barista-stream - hearing ones’ own name or semantic violations - elicited overt gaze-shifts away from the target speaker, above and beyond the baseline level of gaze-shifting (Figure 6).

**Figure 3.**
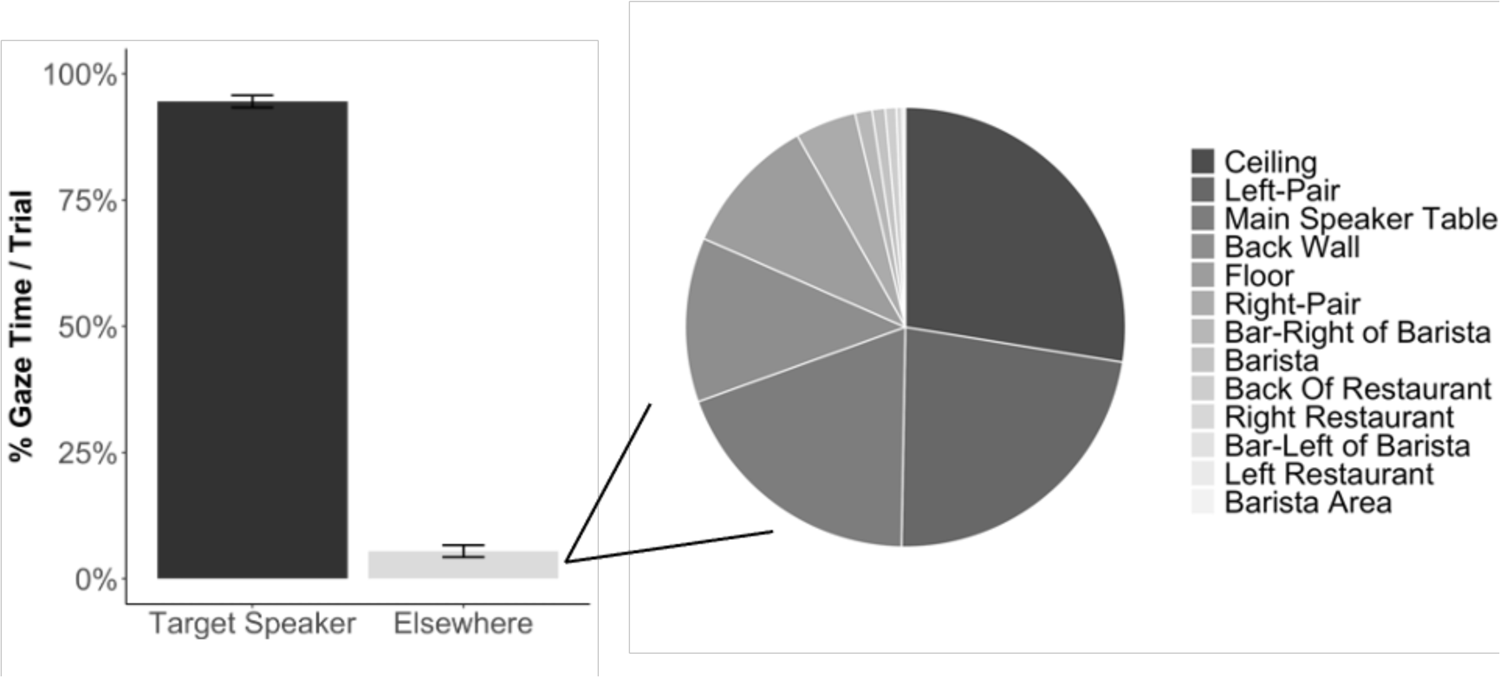
Global distribution of gaze, across participants (n=40). **Left:** Bar graph depicting the percent time of each trial that participants focused their gaze on the target speaker vs. any other parts of the environment. **Right**: Pie chart breaking down the relative time spent looking at ROIs that were <u>not</u> the target-speaker.

### Global gaze-patterns

As expected, participants primarily focused their gaze on the target-speaker, for an average of 94.6 ± 7.4% (std) of each trial (Figure 3, left), confirming that by and large, participants followed instructions and paid attention to the target speaker. When not looking at the target-speaker, the next most popular places to look at were (Figure 3, right): the ceiling, the target speaker’s table and the left-pair of characters (who were more prominent than the right-pair in the default field of view; see Figure 1). The other ROIs, including the Barista and the area around it, drew very few overt saccades (<1% of each trial). The average number of saccades away from the target speaker, at the group level, was 18.64/trial (∼40-second long trials).

However, in looking at individual participants it is interesting to note large differences between them. As illustrated in Figure 4, some participants made frequent saccades away from the target speaker (top example), whereas others remained focused on the target-speaker for the entire trial (bottom example).

**Figure 4:**
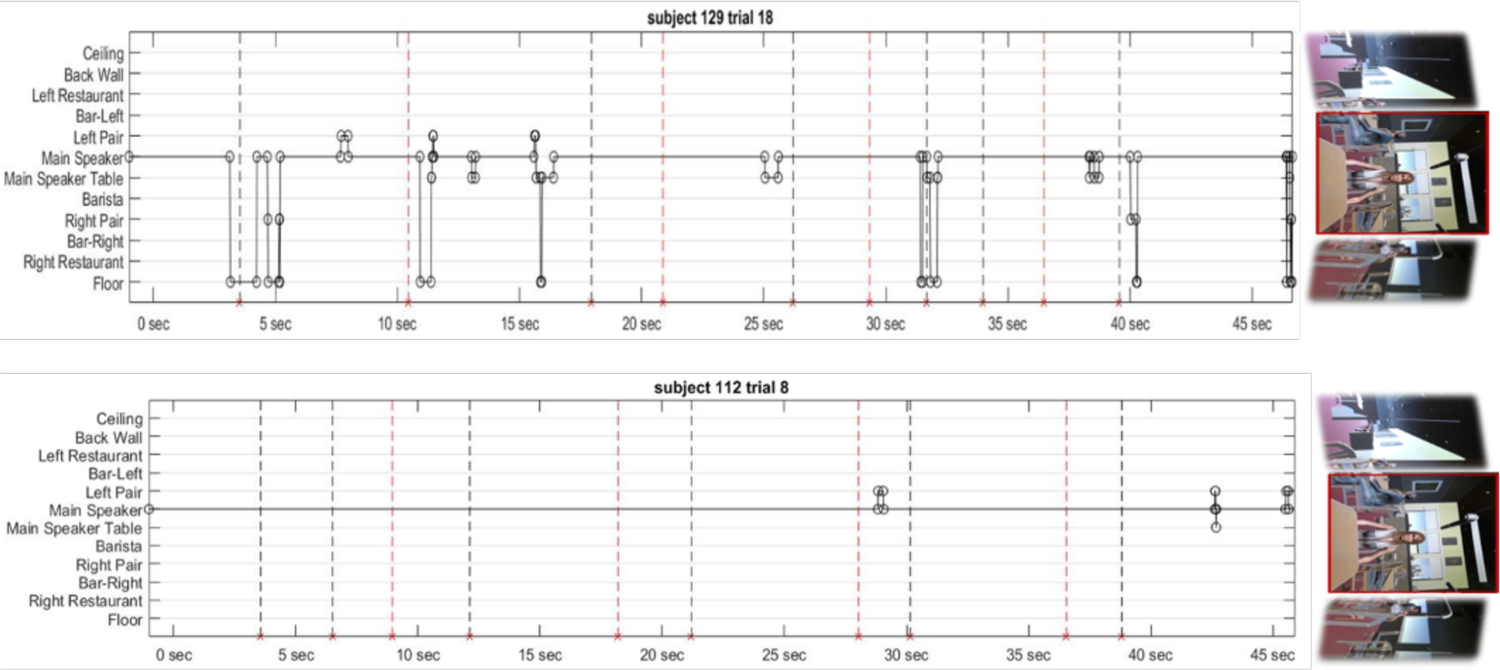
Two examples of different trials from different participants depicting eye movement in relation to targets. Circles represent gaze shift to object. Red dashed lines represent the target probes (both own name and semantic violation), while the black dashed lines represent the respective control probes. Top: represents a trial with a lot of gaze shifts away from the main speaker. Bottom: represents a trial where there weren’t many gaze shifts.

This between-subject variability is portrayed in Figure 5, showing that the average number of saccades per trial *away* from the target-speaker varied between 0-40 (Figure 5A), and the average time spent looking at the target-speaker varied between 100%-60% of each trial (Figure 5B). Not surprisingly, these two measures were significantly correlated (R^2^=0.36, p<0.001; Figure 5C). Interestingly, performance on the word-detection behavioral task was also inversely correlated with the number of saccades performed away from the target-speaker (R^2^=-0.36, p<0.05; Figure 5D), indicating that participants who looked away from the target-speaker more frequently, ultimately detected fewer words. Behavioral performance was not significantly correlated with the overall percent of time spent looking at the target speaker (R^2^=0.05, p=0.147; Figure 5E). Working memory capacity was not significantly correlated with either of our eye-tracking measures (WM vs. frequency of saccades: R^2^=-0.05, p=0.75; Figure 5F, WM vs. percent gaze time at target speaker: R^2^=-0.19, p=0.28).

**Figure 5:**
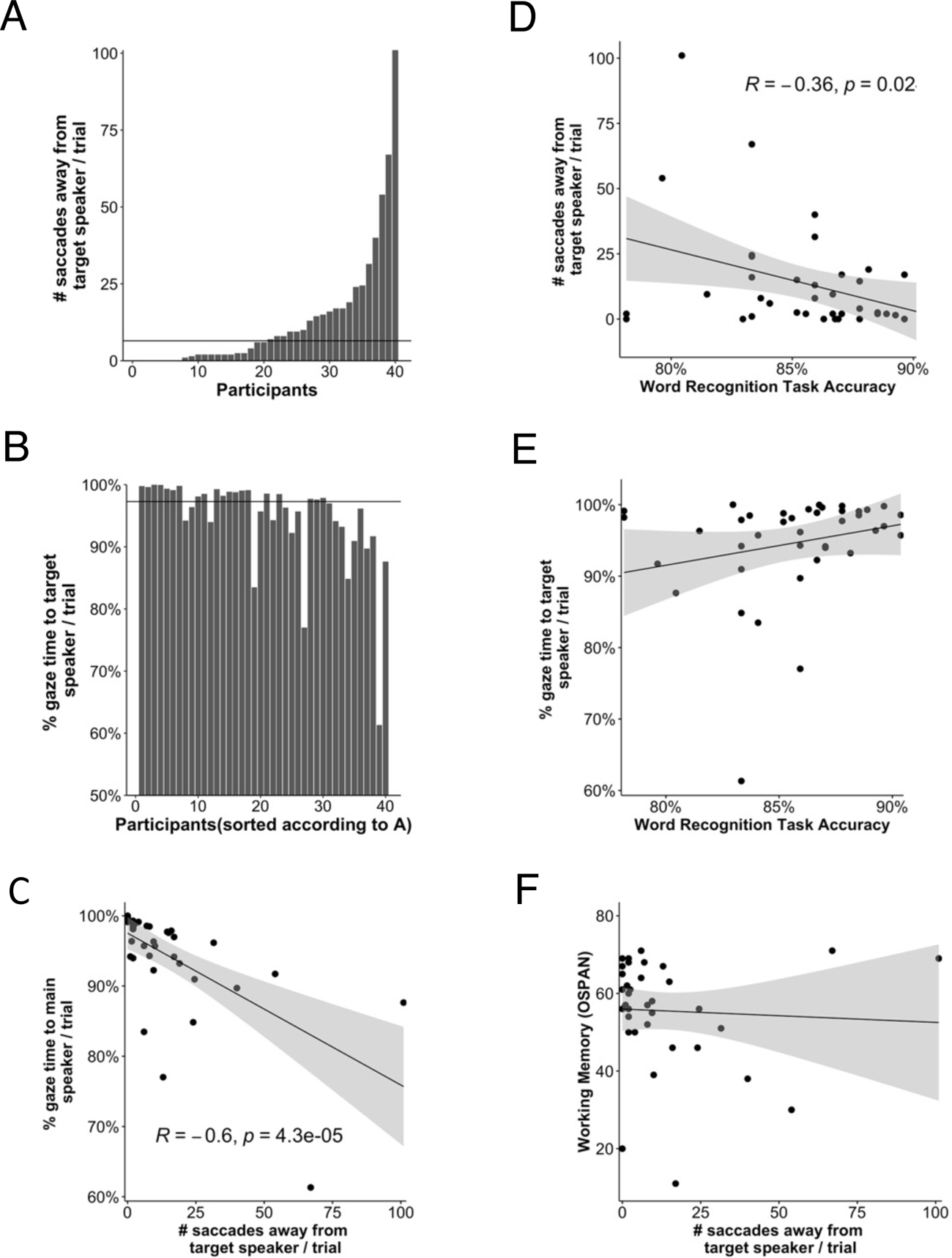
Global eye-gaze patterns across individuals (n=40). **A.** The mean number of saccades performed away from the target speaker per trial, across all participants (sorted). Horizontal line indicates the median across all participants. **B.** The percent of time per trial spent looking at the target-speaker, across all participants (sorted according to A). Horizontal line indicates the median across all participants. **C.** Scatter plot between the two eye movement measures depicted in A & B, which are significantly correlated. **D**. Scatter plot between the behavioral performance in the word recognition task and the number of saccades away from the target speaker, per trial, across participants. Correlation is significant (p=0.02). **E**. Scatter plot between the behavioral performance in the word recognition task and the percent of the trial spent looking at the target speaker. Correlation is non-significant (p=0.147). **F**. Scatter plot of working memory score (OSPAN) task and the number of saccades away from the target-speaker per trial, across participants. Correlation is non-significant (p=0.75).

### Event-related gaze-patterns to probes

Our main interest was in looking at whether the two probe-words – hearing ones’ own name or semantic violations – prompted participants to move their eyes away from the target speaker. To this end, we looked at 2-second epochs following an own-name probe or semantic-violation probe, as well as their respective controls, and counted how many of these epochs contained at least one saccade. When divided by the number of epochs, this constitutes the probability of making a saccade following a probe or a control word. The eye-tracking data for this analysis was only usable in 27 participants. When considering saccades to all ROIs, the Wilcoxon test found no significant difference between the probability to make a saccade after a probe-word vs. a control word (for the name probes, Z=100, p=0.93; for the semantic probes, Z=192, p=0.28; Figure 6A). When limiting the analysis only to saccades from the target-speaker to the barista, we did find a significantly larger probability to make a saccade following a semantic violation vs. its control (Z=26, p<0.05; Figure 6B). However, no significant difference was found between the probability of making a saccade towards the barista after hearing your own-name vs. a control-name (Z=11, p=0.72).

**Figure 6:**
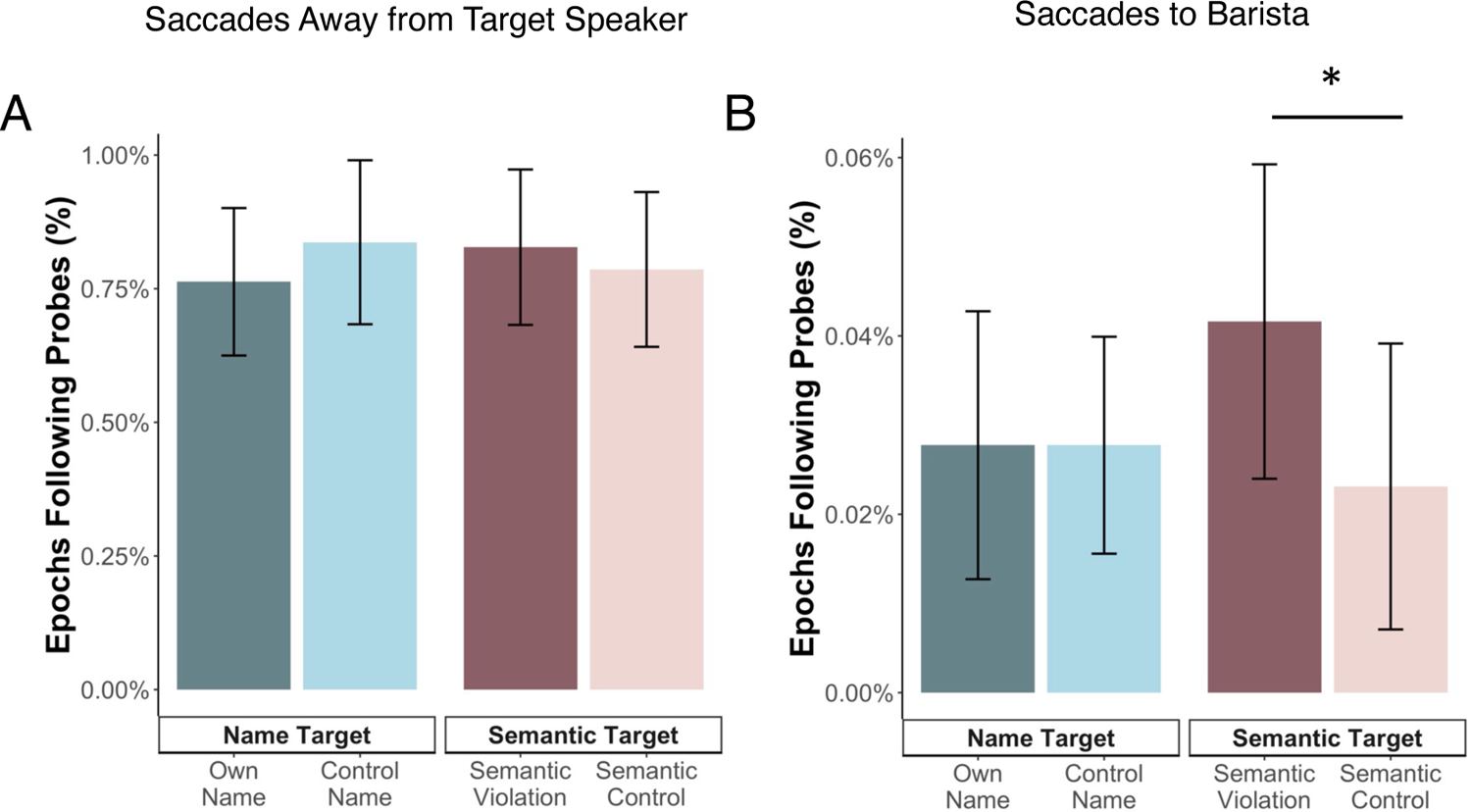
Event-related gaze-shifts following probes (n=27). **A.** Bar graph depicting the percent of 2-second epochs following the presentation of name-probes, semantic-violations, or their respective controls, in which participants performed saccades away from the target speaker (regardless of where the saccade was made to). Error bars indicate SEM. **B.** Bar graph similar to A, but including only saccades performed specifically towards the barista character (located in the back of the café, slightly to the right). Error bars indicate SEM. Asterisk * represent significant difference at a level of p<=0.05.

### EEG results

We estimated the RIDE-ERPs in response to each type of probe-word and their respective controls (5 participants were excluded from this analysis, hence data is reported for n=45). Statistical analysis was performed on the RIDE-ERP grand average across participant, an analysis that accounts for latency variability across trials. However, we also present the traditional ERP grand-average (shown in the insets of Figures 7&8) for qualitative comparison of the resulting waveforms.

**Figure 7.**
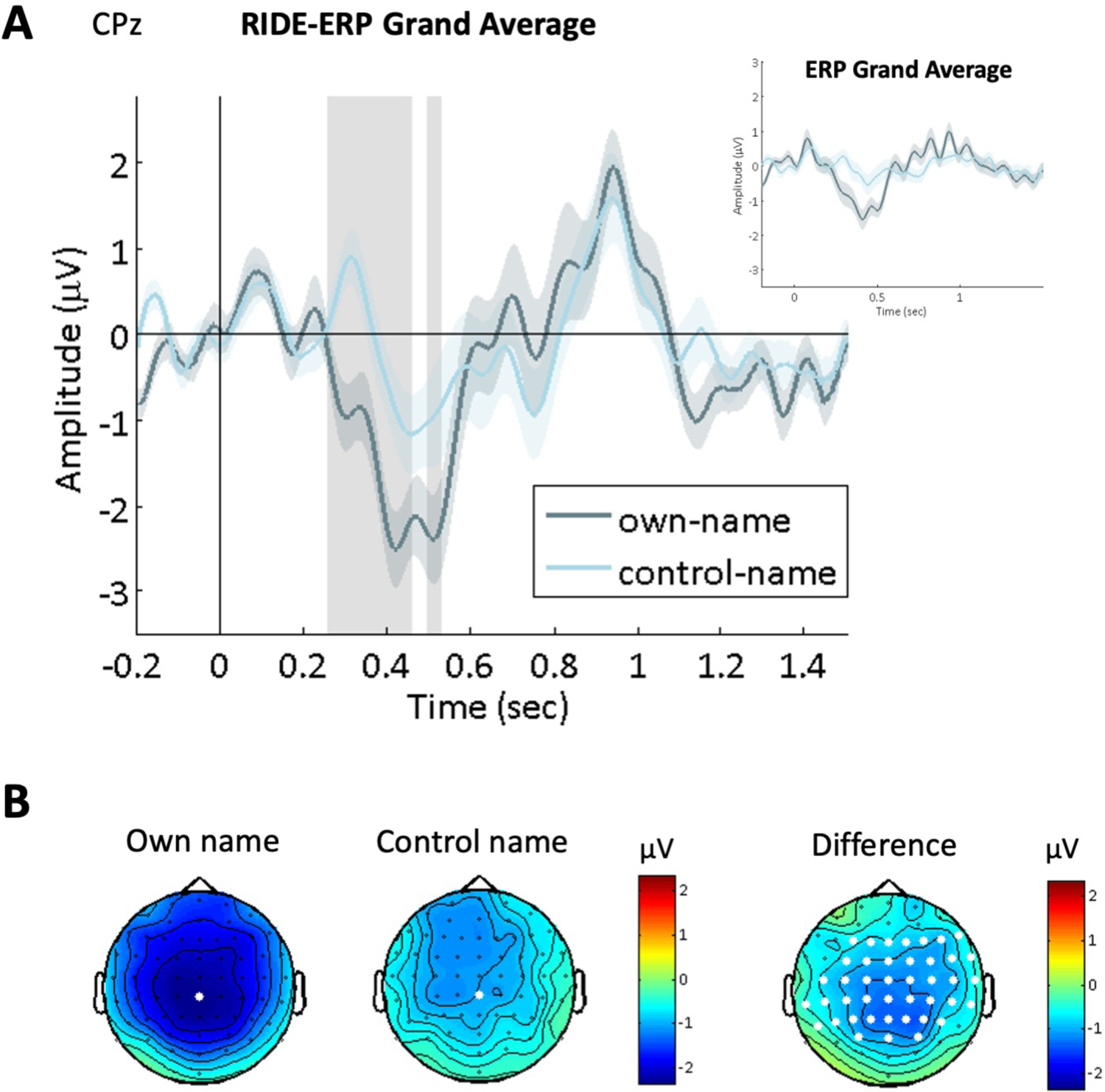
The neural response to own- and control-name probes (n=45). **A.** RIDE-ERP grand-average, shown at electrode CPz (location shown on the topographies in B on the left). Shaded vertical areas indicate the time periods in which significant differences were found between the response to ones’ own vs. control name (p<0.05, TFCE corrected). Inset: Waveform of the classic ERP grand-average, prior to the RIDE analysis. Shaded areas around both waveforms indicate SEM. **B.** Scalp topographies of the responses to the own- and control-name probes (left), and the difference between them (right), in the time window between 400-520 ms. The electrodes where significant differences were found are highlighted in white.

### Detection of own name

Figure 7 shows the comparison between neural responses to hearing ones’ own-name vs. their personal control-name in the barista-stream. Two prominent peaks are observed in the RIDE-ERP grand average – a negative peak between 300-550ms, and a later positive peak between 800-1200ms, both of which were maximal at centro-parietal electrodes. Statistical analysis showed that the mid-latency negative peak was significantly larger in response to ones’ own-name vs. its control, reaching significance primarily between 250**-**460ms (significant electrodes marked in Figure 7B; p<0.025, two-tailed TFCE cluster correction).

### Detection of semantic violation

Figure 8 shows the comparison between neural responses to words that create a semantic violation vs. well-formed sentences in the barista stream. The RIDE-ERP grand average to these words showed a series of three peaks – a positive peak between 250-450ms, a negative mid-latency peak between 600-800ms, and a later positive peak between 900-1200ms. Statistical analysis indicated that the positive response between 200-420ms was significantly larger in response to semantic violations vs. their controls (significant electrodes marked in Figure 8B; p<0.025, two-tailed TFCE cluster correction).

**Figure 8.**
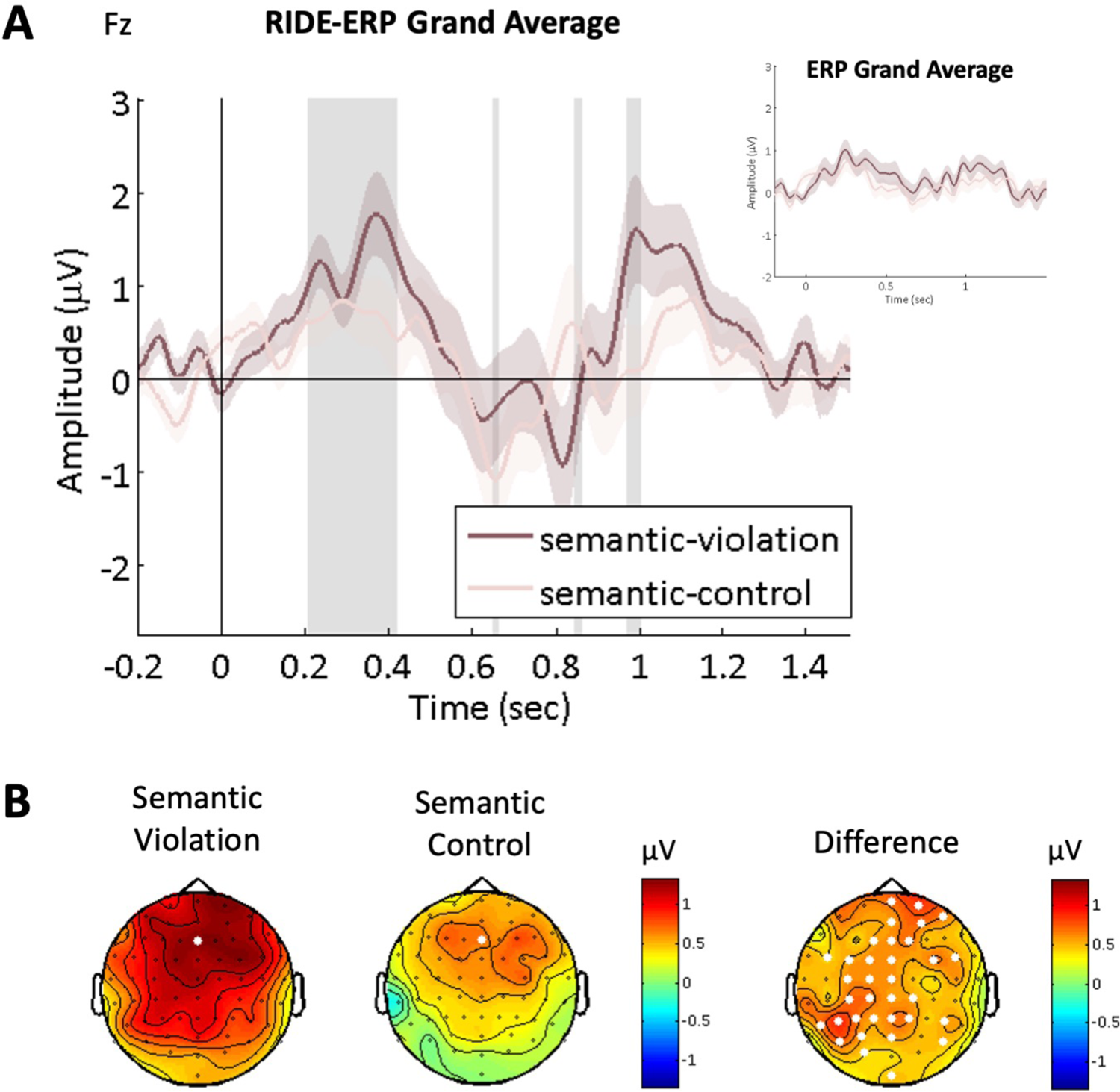
The neural response to semantic-violations vs. control probes (n=45). **A.** RIDE-ERP grand-average, shown at electrode Fz (location shown on the topographies in B on the left). Shaded vertical areas indicate the time periods in which significant difference was found between the response to semantic violations vs. their controls (p<0.05, TFCE corrected). Inset: Waveform of the classic ERP grand-average, prior to the RIDE analysis. Shaded areas around both waveforms indicate SEM. **B.** Scalp topographies of the responses to the own- and control-name probes (left), and the difference between them (right), in the time window between 210-420 ms. The electrodes where significant differences were found are highlighted in white.

### GSR results

Figure 9 shows the comparisons between the grand-averaged GSR responses to hearing one’s own-name and semantic-violation vs. their respective controls. Both waveforms show a wide positive peak in response to the probe-words, but not to their controls. Statistical comparison of the mean response in a 2-4sec time window following each probe showed significantly higher GSR responses to hearing ones’ own name vs. a control-name (p=0.0207, one-tailed t-test), as well as a higher response to semantic violations vs. their control (p=0.0364, one-tailed).

**Figure 9:**
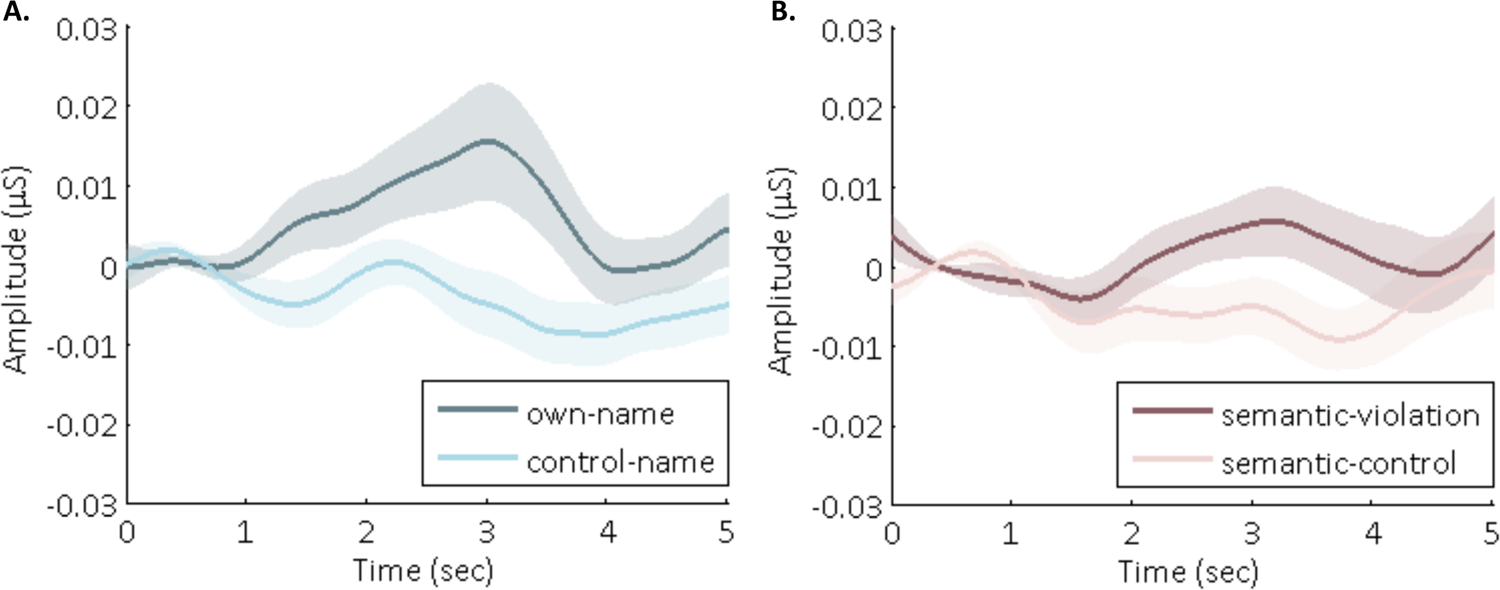
GSR grand-averaged responses to **A.** hearing ones’ own-name vs. control-name, and **B.** hearing the semantic-violation probes vs. control words. (n=50)

## Discussion

This study introduces a novel and highly ecological experimental approach for studying how individuals manage their attention in realistic “cocktail-party” type situations. We pursued two main lines of investigation: First, we examined the pattern of spontaneous eye-gaze dynamics as participants attended to a target-speaker while immersed in the rich and realistic audiovisual setting of a virtual café. Second, we tested whether there is evidence for incidental detection of semantically and personally salient words in task-irrelevant background speech, under these ecologically valid conditions. Results from both lines of investigation reveal important insights about how attention is allocated in busy natural settings, highlight the importance of considering individual differences in studies of attention, and raise questions regarding the underlying assumptions of some more traditional selective attention studies.

### Spontaneous gaze-dynamics

As expected, participants primarily focused their gaze on the target-speaker, and accordingly displayed good behavioral recall of words from her speech. However, a closer look at the gaze-patterns of individual participants revealed a slightly more complex pattern. While some participants looked almost exclusively at the target-speaker and their gaze did not waver, others performed a substantial number of saccades around other locations in the café. Perhaps not surprisingly, the frequency of saccades was negatively correlated with performance, and participants who looked around more often ultimately recalled less of the target speech. This is in line with results from a previous experiment of selective attention to speech in a virtual café (Shavit-Cohen and Zion Golumbic, 2019), where we found large inter-subject variability alongside high intra-subject consistency in the tendency to either stay focused or look around the room. The latter behavior presumably reflects an instinct to occasionally scan the environment for potentially interesting information or a personal inclination towards frequent attention-shifts (Oberfeld and Klöckner-Nowotny, 2016; Ziegler et al., 2018). We tested a specific proposed hypothesis that tendency to monitor task-irrelevant portions of the environment is related to differences in working-memory capacity (Conway et al., 2001; Beaman, 2004; Colflesh and Conway, 2007; Gazzaley and Nobre, 2012; Cak et al., 2020; Lambez et al., 2020). However, our data to not support this hypothesis, as neither the gaze-dynamics nor behavioral performance were correlated with WMC (as operationalized here). Other hypotheses regarding the factors contributing to these behaviors, to be tested in future research, include traits such as personality traits, perceptual curiosity, or tendencies of hyperactivity or inattention (Hoppe et al., 2018; Bendall et al., 2021; Maron et al., 2021; Zangrossi et al., 2021; Khatri et al., 2022; Mauriello et al., 2022; Stokes et al., 2022). Regardless of the source of this inter-subject variability, this pattern reminds us that although selective attention studies usually assume that participants allocate their attention ‘as instructed’, in practice overt attention (as well as covert attention) may fluctuate and shift spontaneously, in manners that the experimenters cannot always monitor. Taking the potentially dynamic nature of attention into account is crucial for advancing attention research into more realistic and ecological realms.

### Incidental detection of salient words in task-irrelevant speech

Our results demonstrate that even as listeners engaged in selective attention towards the target-speaker, this stimulus was not processed exclusively, and some resources were also devoted to processing the competing barista speech. We found clear neural and physiological responses to **both** probes – hearing ones’ own name and semantic violations - in the task-irrelevant barista speech, supporting the notion that task-irrelevant speech is not merely encoded at the acoustic level, but is also processed for linguistic content and semantic salience. As discussed above, the degree to which task-irrelevant speech is processed has been a topic of heated debate, and conflicting results have been reported over the years (Paavilainen, 2013; Ding et al., 2018; Har-shai Yahav and Zion Golumbic, 2021; Röer and Cowan, 2021), though mostly using highly artificial tasks and stimuli. The current study substantially broadens this conversation, by showing evidence for the incidental detection of words in task-irrelevant speech, in an ecologically valid context simulating situations and stimuli that people may actually encounter in real-life.

### Own-name detection

The special behavioral and neural sensitivity to hearing ones’ name has been established during passive listening (mostly to lists of words) or when participants engage in a visual task (Berlad and Pratt, 1995; Folmer and Yingling, 1997; Müller and Kutas, 1997; Perrin et al., 1999, 2005; Holeckova et al., 2006; Höller et al., 2011; Tamura et al., 2012; Tateuchi et al., 2012; Eichenlaub et al., 2012; Röer et al., 2013; Del Giudice et al., 2014; Ljungberg et al., 2014; Lechinger et al., 2016; Jijomon and Vinod, 2021). This own-name advantage has been hypothesized to reflect a special status of self-representation (Gallagher, 2000; Kricher and David, 2003; Perrin et al., 2005; Röer et al., 2013), possibly related to an evolutionary survival mechanism (Müller and Kutas, 1997). However, whether or not people also detect their name when they actively pay attention to another speech stream, has yielded conflicting results (Cherry, 1953; Moray, 1959; Wood and Cowan, 1995; Naveh-Benjamin et al., 2014; Holtze et al., 2021). Part of the difficulty in ascertaining the extent to which people detect their name in task-irrelevant speech has been methodological: This can only be assessed based on subjective recall at the end of the experiment (Moray, 1959; Conway et al., 2001; Röer and Cowan, 2021) or by indirect assessment of its interference with main task performance (Wood and Cowan, 1995; Röer et al., 2013; Ljungberg et al., 2014; Naveh-Benjamin et al., 2014). The combined recording of neural (EEG) and physiological (GSR) responses in the current study, allowed us to directly evaluate whether participants detected their own name, without the need to resort to indirect or subjective reports.

Using the RIDE-analysis, to overcome latency differences in neural responses to stimuli of different lengths, we found that hearing ones’ own name in the barista-stream elicited an increased negative response in the RIDE-ERP around 400ms. While we acknowledge that the RIDE-ERP may not be directly comparable to traditional ERPs, the time-scale, polarity and scalp-distribution of the own-name effect found here, is consistent with the well-established N400 ERP component, which has been shown to be enhanced in response to hearing ones’ name in less ecological paradigms (Müller and Kutas, 1997; Eichenlaub et al., 2012; Tamura et al., 2012). Other ERP studies looking at response to hearing ones’ name found that it led to enhanced P300 responses, that is traditionally associated with target-detection rather than semantic processing (Berlad and Pratt, 1995). Notably, a recent study by Holtze *et al*. (2021), found that hearing ones’ name in a task-irrelevant speech stream elicited an increased P300 response and as well as a transient increase in speech-tracking of that speaker, a pattern consistent with a transient shift of attention towards task-irrelevant speech, prompted by hearing your name. The many operational differences in the design and stimuli of the work by Holtze *et al*. (2021) vs. the current study can easily account for the differences in the specific neural manifestation of the own-name advantage. However, importantly, they both converge in demonstrating that even when your name is uttered by a speaker that is outside of your primary focus of attention, it ‘pops out’ and is detected by the brain.

This neural response was also accompanied by physiological response of increased arousal, which manifest as a significantly increased galvanic-skin-response (GSR) to hearing one’s own name in the barista-speech. This is in line with previous findings showing heightened GSR in response to familiar or personally-relevant stimuli (Gronau et al., 2003) as well as response to ones’ own name when it is task-relevant (Pinto et al., 2022). However, to the best of our knowledge this has not been shown previously for names presented in task-irrelevant speech, serving as converging evidence for its incidental detection and unique perceptual status.

### Detection of semantic violations

Besides the singular example of detecting ones’ own name, including semantic violations in the barista-speech allowed us to expand the question of how task-irrelevant speech is processed. The basic rationale is that finding an indication for detection of semantic violations, implies that the entire speech was monitored for its semantic content, in line with late-selection models of attention (Parmentier et al., 2018). As in the case of detecting ones’ own name in task-irrelevant speech, behavioral studies offer conflicting results regarding the degree to which the presence of semantic violations in task irrelevant speech interfere with performance of a main task (Bentin, Kutas and Hillyard, 1995; Röer *et al*., 2019, 2021;, vs. Deacon, 2000; Aydelott, Jamaluddin and Nixon Pearce, 2015; Röer *et al*., 2017; Parmentier *et al*., 2020; Röer and Cowan, 2021). Hence here, too, monitoring neural and physiological responses to semantic violations in the barista-stream provides a more direct, and ecologically relevant, metric.

Semantic violations are known to elicit an enhanced N400 neural response when present in task-relevant stimuli, effects shown primarily for text (Marta and Hillyard, 1980; Kutas and Federmeier, 2011; Luka and Van Petten, 2014) but also for speech (Holcomb and Neville, 1990; Hahne and Friederici, 2002; Balconi and Pozzoli, 2005). The few studies that have looked at whether similar effects are found for task-irrelevant speech did not find evidence for an enhanced N400 response (Bentin et al., 1995; Kanerva, 2021). Here too, our RIDE-ERP analysis of neural responses to semantic violations did not show a pattern consistent with an N400, however we did find an increased positive response around 300ms, which is more akin to the P300 component, that is associated with target-detection and the capture of exogenous attention (Cid-Fernández et al., 2014; Pinheiro et al., 2017; Masson and Bidet-Caulet, 2019).

Why did we find that semantic violations elicited a P300-like response whereas hearing ones’ name elicited a N400-like response? At this point, we cannot offer a full explanation for this. Additional replications and extensions of this line of research are required to fully characterize the nature of the neural responses to salient words in task-irrelevant speech. Moreover, it is important to acknowledge that the reported effect for semantic-violations was notably weaker than that effect found for hearing ones’ name, and was also more susceptible to parameter choices in the RIDE analysis (see supplementary material; probably due to the more variable nature of the stimuli, which were less uniform than repetitions of one’s name). Bearing these limitations in mind, the converging EEG and GSR results, showing responses for both the own name and semantic violation manipulations, provide strong indications the task-irrelevant barista-speech was monitored by the listeners, and that they were able to glean salient semantic information from it. This converges with the results of several recent studies, showing that listeners are actually capable of following the content of two simultaneous speakers, at least in contexts with moderate acoustic load (Agmon et al., 2021; Har-shai Yahav and Zion Golumbic, 2021; Kaufman and Zion Golumbic, 2022; Pinto et al., 2022).

### Incidental detection not accompanied by overt gaze-shifts

Interestingly, in the current dataset, hearing ones’ name in the barista-stream was not accompanied by overt gaze-shifts towards the barista character, and semantic violations elicited gaze-shifts in an extremely small proportion of instances. This may be because detection of the name remains represented at a subthreshold – perhaps pre-conscious level (Dehaene et al., 2006), or because social conditioning of social appropriateness inhibits listeners from performing overt gaze-shifts (Richardson and Gobel, 2015). This pattern invites further exploration of the relationship between the internal detection-processes indexed by the neural and physiological responses, and the elicitation of overt gaze-shift.

## Conclusion

This study is part of a larger endeavor to study how humans deal with the perceptual complexity and cognitive challenges of real-life environment. The use of an immersive virtual-reality experimental platform, where participants contend with the type of audio-visual stimuli they might encounter in real life, brings us closer to this goal and affords insights into how individuals navigate these complexities. The combined neural, ocular, and physiological data reported here demonstrate the non-exclusive nature of real-life attention, whereby even if one stimulus is of primary interest to an individual, they also scan the environment occasionally and pick up words from other speech stimuli around them. Despite proposed ‘bottlenecks’ for linguistic processing of multiple speech stimuli, our data suggest that sufficient resources are available to allow processing of a secondary speech stream, at least under the ecologically-valid levels of perceptual and cognitive load tested here. The dynamic perspective of real-life attention implied by the current results is in line with the notions of active sensing and attentional sampling (Morillon et al., 2015; Leszczynski and Schroeder, 2019; Fiebelkorn and Kastner, 2020; Plöchl et al., 2022) and likely carries ecological benefits, allowing individuals to notice and react to important events around them, rather than approaching the world with ‘tunnel vision’ where all available processing resources are allocated to only one stimulus of interest.

## Acknowledgements

This work was supported by the Israel Science Foundation grant # 2339/20 (to EZG) and the Israel Ministry of Science grant # 88962 (to EZG). We would like to thank Dr. Maya Kaufman for consulting on stimulus design and Ms. Orel Levi for assistance in data collection.

