## Supplemental Figures for "Covert detection of own-name and semantic violations in task-irrelevant speech, in a realistic Virtual Café"

Supplementary Material

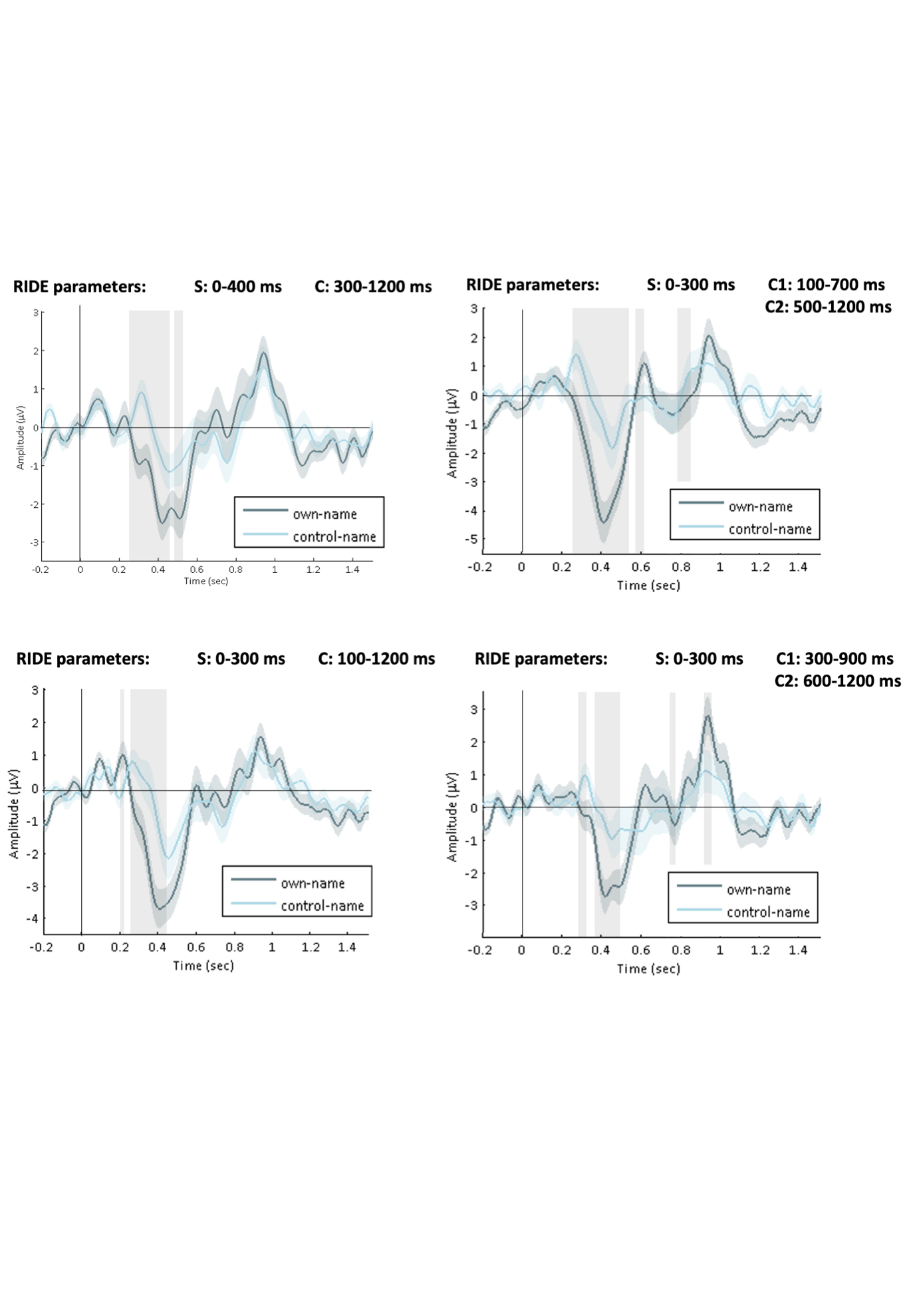

**Figure S1**. Grand average RIDE-ERP to ones’ own name vs. its control from electrode CPz, derived using different parameter choices (Ouyang *et al.*, 2011) (http://cns.hkbu.edu.hk/RIDE.htm). Right column: RIDE models that included a single C component. Left column: RIDE models that included two C components (C1 and C2). The time-windows defined for each component are stated above each figure. Shaded areas around each waveform indicate the standard error of the mean. Shaded vertical areas indicate the time periods in which significant difference were found between the two response (p<0.05, TFCE corrected). The modulation of a negative peak around 400 ms in response to hearing ones’ own name was significant in all iterations of this analysis, supporting its robustness. Data from top-right panel is reported in the main paper.

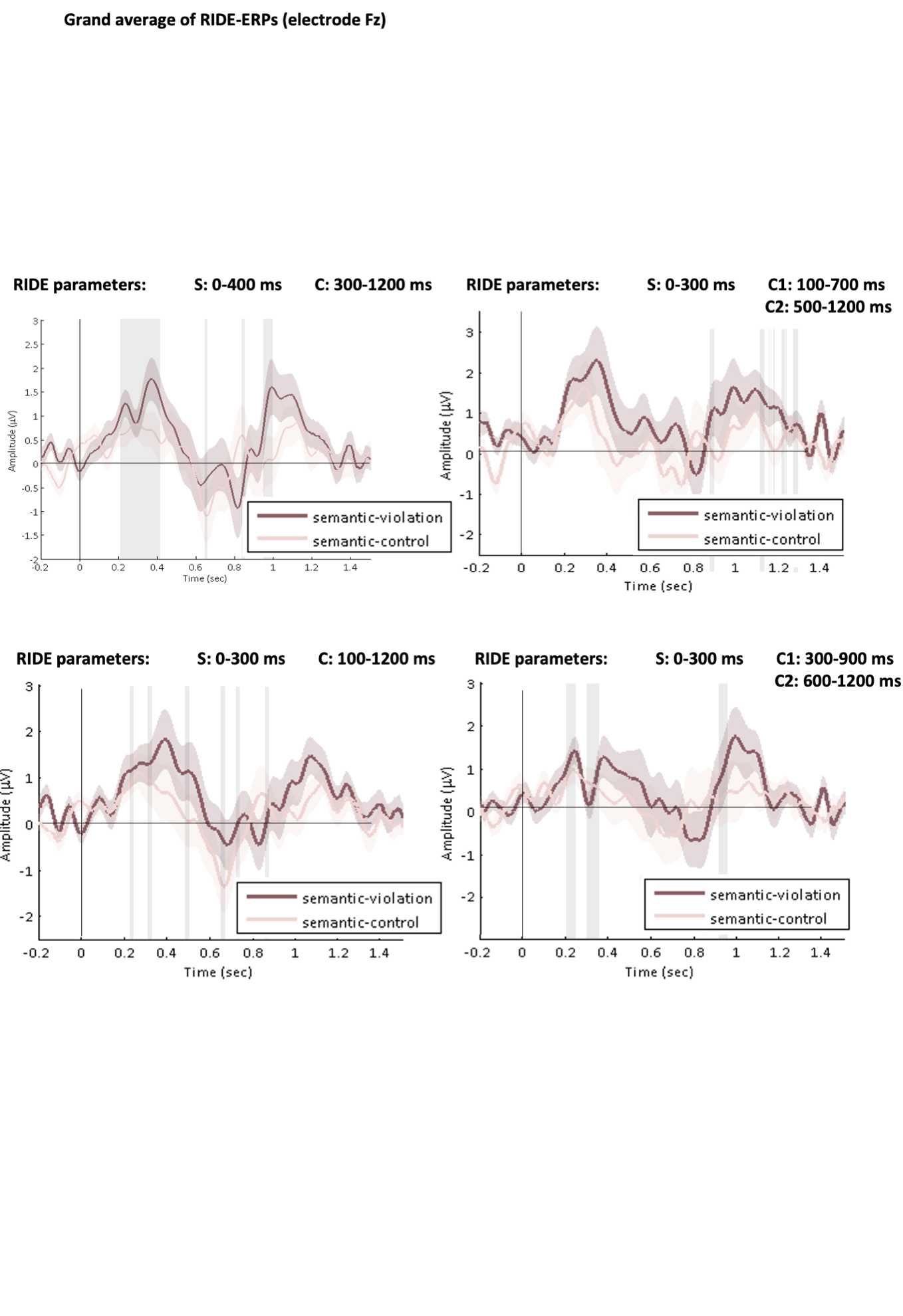

**Figure S2**. Grand average RIDE-ERP to semantic violations vs. its control from electrode Fz, derived using different parameter choices (Ouyang *et al.*, 2011) (http://cns.hkbu.edu.hk/RIDE.htm). Right column: RIDE models that included a single C component. Left column: RIDE models that included two C components (C1 and C2). The time-windows defined for each component are stated above each figure. Shaded areas around each waveform indicate the standard error of the mean. Shaded vertical areas indicate the time periods in which significant difference were found between the two response (p<0.05, TFCE corrected). The modulation of a positive peak between 200-400 ms in response to semantic violations was more susceptible to parameter choices in the RIDE analysis, and was not significant in all iterations of this analysis (although the trend is observed in most parameter choices). This may be due to the more variable nature of the stimuli averaged here (which were no simply repetitions of the same word, as in the own-name case), and we would recommend replication attempts of these results in future research. Data from top-right panel is reported in the main paper.
